# Congruent brain signatures specific to phonology in fronto-temporal cortex during language production and understanding

**DOI:** 10.1101/2024.09.22.614347

**Authors:** Xenia Dmitrieva, Jean-Luc Anton, Elin Runnqvist, Amie Fairs, Bissera Ivanova, Julien Sein, Bruno Nazarian, Sophie Dufour, Friedemann Pulvermuller, Kristof Strijkers

## Abstract

In this fMRI study we investigated whether language production and understanding recruit the same phoneme-specific networks. We did so by comparing the brain’s response to different phoneme categories in minimal pairs: Bilabial-initial words *(e.g., ‘**m**onkey’)* were contrasted to alveolar-initial words *(e.g., ‘**d**onkey’)* in 37 participants performing both language production and comprehension tasks. Region-of-Interest analyses showed that the same sensorimotor networks were activated across the language modalities. In motor regions, word production and comprehension elicited the same phoneme-specific topographical activity patterns, with stronger haemodynamic activations for alveolar-initial words in the tongue cortex and stronger activations for bilabial-initial words in the lip cortex. In the posterior and middle superior temporal cortex, production and comprehension likewise resulted in similar activity patterns, with enhanced activations to alveolar-compared to bilabial-initial words. These results disagree with the classical separation between language production and understanding in neurobiological models of language, and instead advocate for a cortical organization where the same phoneme-specific acoustic-and-articulatory representations carry production and comprehension.

## Introduction

The production and comprehension of language must be continuously coordinated in dialogue and conversation ^(1,2)^. Therefore, understanding how production and comprehension interact with each other is important for cognitive and neurobiological models of language. Nevertheless, historically, the language modalities have been studied separately. This was in part due to observations with aphasic patients, who were once classified in ‘sensory’ and ‘motor aphasias’, suggesting that each language modality relies on distinct processing pathways in the cerebral cortex ^(3,4)^; an observation which seemed to receive further support from neuroimaging work with healthy participants ^(5)^. However, several findings cast doubt on a neat separation of production and comprehension mechanisms in encapsulated modules: While ‘sensory aphasia’ is well-known to be characterized not only by comprehension deficits, but by production deficits too ^(6,7)^, even aphasias once believed to be entirely ‘motor’ come with well-characterized comprehension deficits ^(8-10)^. Similarly, fine-grained analyses of language production and comprehension behavior in healthy language users suggests a closer relationship between the two systems than previously assumed ^(1,11-13)^. However, neurometabolic and neurophysiological imagining studies are necessary for more precisely defining that relationship and, potentially, the degree of overlap between the brain’s production and comprehension networks.

In recent years, interest for cross-modal language research has notably increased and several neuroimaging studies have contrasted production and comprehension for conversational dynamics and sentence processing. This intriguing work demonstrated strong neural overlap across the language modalities, suggesting brain systems for syntax, message-level semantics and story-level content that overlap not only in the classic left-hemispheric frontotemporal language areas but also include a range of additional left- and right-hemispheric loci ^(14-19; but see: 20)^. We note however, that most of these studies engaged their participants in tasks that exceed basic language comprehension or production per se, and involve a range of complex cognitive processes such as theory-of-mind, indirect inferences of utterance meaning, prediction of upcoming phrasal content, memory recall, attention and so forth. While we certainly agree that all of these processes are necessarily paired with the use and understanding of the basic building blocks of language, nevertheless, when it comes to drawing strong conclusions on any shared brain mechanisms for language production and comprehension, it is imperative to distinguish specific language representations from any of the aforementioned accompanying and higher-level cognitive operations triggered by the complex linguistic tasks. Therefore, here we investigate the neural overlap between the language modalities with functional magnetic resonance imaging (fMRI) using well-known, basic tasks at the word-level and focus on the question whether phoneme-specific brain signatures are present in both the production and comprehension of speech.

Focusing on phonological processing at the word-level is interesting since current brain language models strongly differ with regard to their assumptions about the separation or integration of phoneme representations in word production versus comprehension. Broadly speaking, two classes of models can be contrasted: According to Partial Separation Models (PSM), both frontal and temporal brain regions are recruited for phonological and word form processing in language production, whereas only temporal regions are needed for speech comprehension ^(21-26)^. An alternative proposal is a class of models we label Integration Models (IM), according to which both language production and comprehension recruit the same frontal and temporal regions during phoneme and word form processing ^(27-33)^.

At present, there are only two studies we know of that have directly compared phonological processing between the production and comprehension of words within the same participants. In Okada & Hickok ^(34)^ fMRI data demonstrated neural overlap for word production and comprehension in the posterior superior temporal gyrus and sulcus (pSTG and pSTS), regions which are often linked to phonological processing ^(21)^. However, in Okada & Hickok, no specific phonological manipulation was targeted, which makes inferences about phonology indirect; speech production was assessed covertly, which has been criticized to be distinct from natural overt production ^(35)^; furthermore, the number of tested subjects was rather low (10 participants). More recently, Fairs and colleagues ^(36)^ contrasted with EEG the passive perception and overt production of words varying in phonotactic frequency (i.e., how often phonemes in a word co-occur ^(37)^). The authors observed early ERP modulations elicited by phonotactic frequency in both production and comprehension, making them conclude that the same rapid parallel brain dynamics underpin both modalities. Nevertheless, overlapping temporal dynamics between the production and comprehension of words does not mean that phonological processing also overlaps in neural representations. In sum, the few neurophysiological data available thus far supports the idea of some cross-modal overlap in a word’s phonology, but these data need to be extended. The current study aims at doing so by comparing with fMRI the overt production and passive perception of words within the same participants, and utilizing a specific phonological manipulation which allows us to directly assess whether **(1)** topography-specific networks in frontal and temporal cortex underpin word phonology, and **(2)** whether such phonological networks are non-overlapping in production and comprehension as predicted by PSM or whether they are the same across the language modalities as predicted by IM.

To tap into phonology, we compare the mapping of word-initial phonemes in minimal pairs, which are words that differ only in a single sound. Concretely, we contrast minimal pairs with either bilabial- or alveolar-initial word phonemes (e.g., */**<u>b</u>**allon/* vs */**t**alon/* in French; *English translation: ‘ball’ vs. ‘heel’*). Using this contrast is interesting because it concerns an unambiguous phonological manipulation which, unlike certain formal variables like word length or neighborhood density, is not correlated with other stages of linguistic processing ^(38)^. Also, the manipulation of bilabial and alveolar phonemes has already been successfully applied to both the production ^(39-42)^ and perception of speech ^(43-47)^. This makes it an ideal contrast to utilize in a study that aims to directly compare both modalities. Of particular relevance are the studies by Pulvermuller and colleagues in perception ^(44)^ and Strijkers and colleagues in production ^(42)^. In an fMRI study, Pulvermuller et al. found that listening to bilabial and alveolar speech syllables (e.g., */ba/* vs. */ta/*) activated the motor cortex in a topography-specific manner. Bilabial speech sounds activated more strongly those regions responsible for moving our lips and alveolar speech sounds activated more strongly the motor cortex responsible for moving our tongue. Using magnetoencephalography (MEG), Strijkers et al. extended this finding to speech production with real words (e.g., */Donkey/* vs */Monkey/*), observing topography-specific activation of the motor cortex within 200 ms after presentation of to-be-uttered object names. In addition, both studies also reported phoneme-specific activations in the pSTG of place of articulation, suggesting a correspondence in fronto-temporal networks sensitive to phoneme processing as hypothesized by IM.

Nevertheless, the critical test that explores within the same brains whether the phonology-sensitive cortical regions activated in response to the production of words are indeed identical to those activated for the comprehension of words, is still lacking. Furthermore, several studies using electrocorticography (ECoG) question a role for place of articulation during speech perception ^(48-50)^: For instance, Cheung et al. ^(50)^ found that while the articulation of speech sounds resulted in motor somatotopy, the perception of speech sounds was not sensitive to place of articulation, but triggered acoustic activations in the motor cortex similar as to the neural responses in the auditory cortex. However, since speech perception involved a phoneme matching task and only meaningless syllables were used, it does not exclude motor somatotopy under normal listening conditions of meaningful language ^(51)^. In the current study, to provide a direct test of the phonology-sensitive brain regions involved in the production and comprehension of meaningful language, we record fMRI in 37 participants who do both a standard production (object naming) and perception task (passive listening) for the same minimal pairs which only differed in place of articulation (***Fig. 1***). With this design we will verify whether phoneme representations of words manifest in the brain in a topography-specific manner in the frontal cortex and possibly in the temporal cortex, and whether such topography-specific phonological network is shared between language production and comprehension. If the predictions of PSM are right, we should observe a modality-specific activation pattern ^(21-27)^: Whereas for the production task both STG and the motor cortex should become activated alike for words with alveolar and bilabial speech sounds, the perception task should spark the temporal cortex exclusively (***Fig. 1***). Note furthermore that some accounts within the class of PSM postulate that also temporal cortex activations will be largely non-overlapping between the perception and production of phonemes ^(21,23,25)^. In contrast, the IM predict topography-specific brain responses in the motor cortex reflecting the somatotopy of the articulators most relevant for pronouncing the critical phonemes, namely the lips for bilabials and the tongue for alveolars, in both language production and comprehension ^(27-33)^. Likewise, also in the STG the phoneme manipulation should trigger the same brain regions when producing or comprehending words, possibly also generating topography-specific responses as in the motor cortex as suggested by some IM ^(28,31)^ (***Fig. 1***).

**Fig. 1.**
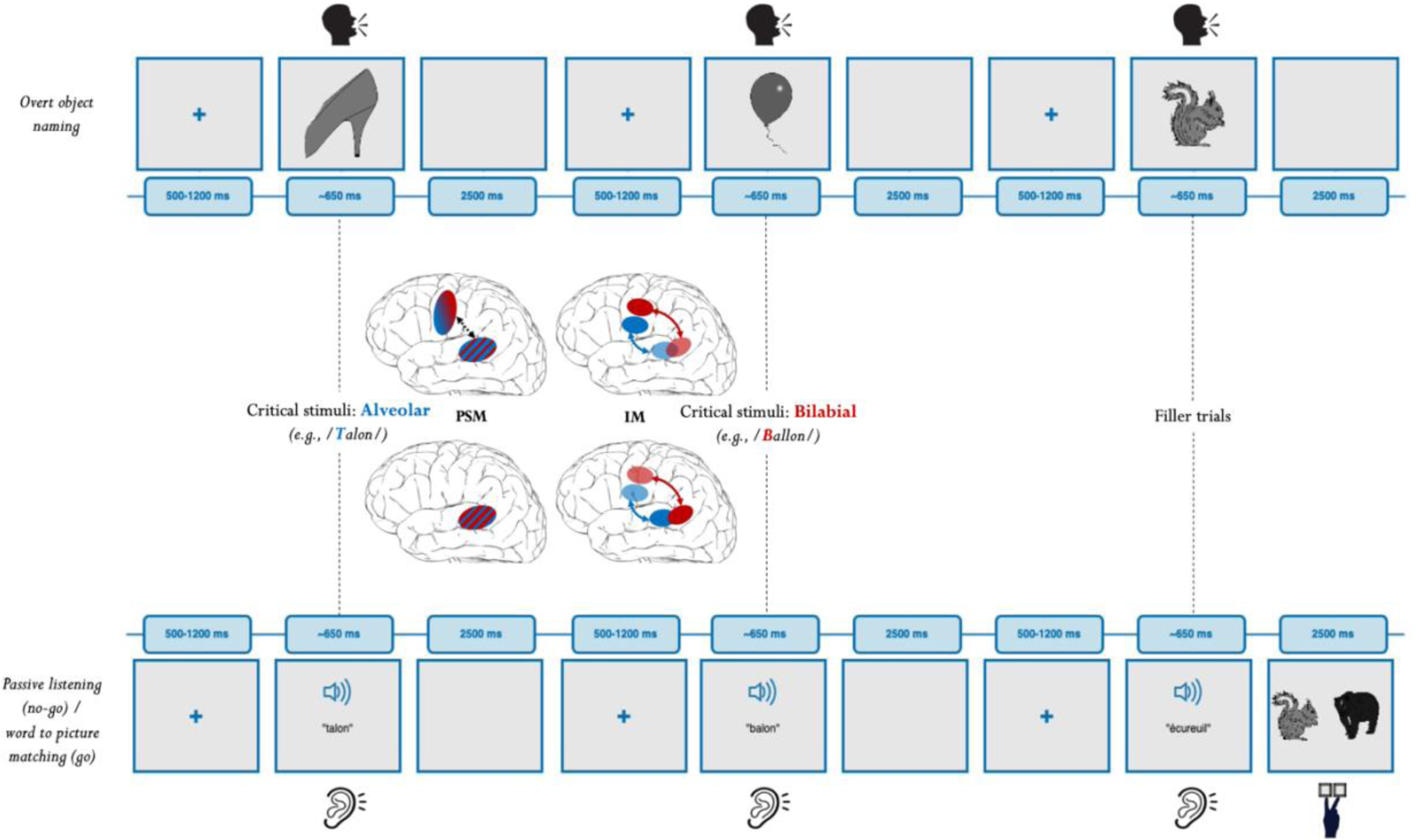
Design and hypotheses of the experiment. The upper panel depicts the trial structure for the language production task (object naming) and the lower panel for the language comprehension task (passive listening). The same lexical units are targeted in both conditions, naming of object pictures by producing minimal pairs of alveolar-initial words (blue) versus bilabial-initial words (red) and listening to the same spoken word forms. The middle panel depicts the hypotheses for this alveolar-bilabial contrast between the language modalities for the Partial Separation Model (PSM) and Integration Model (IM): PSM predict both temporal and motor activation in production, but only temporal activation in perception (which furthermore could be different from the temporal activation in production). IM predict the same fronto-temporal network in production and perception, with different topographically-specific activation patterns in the motor cortex for otherwise well-matched alveolar- and bilabial-initial words (and possibly a somewhat similar topographical response in temporal cortex).

## Results

For clarity, here we will only report the results for the interactions of relevance for our hypotheses. The detailed results of all ANOVAs can be consulted in *the online repository:* https://osf.io/bhfdw/.

Starting with the 4-way ANOVA, which includes all variables tested in this study, namely ‘Phoneme’ (bilabial vs. alveolar) x ‘Area’ (motor cortex vs. superior temporal cortex) x ‘Region of Interest (ROI)’ (anterior (tongue-related) vs posterior (lip-related)) x ‘Modality’ (production vs. comprehension), we observed significant 3-way interactions between ‘Phoneme’, ‘Area’, and ‘ROI’ (*F(1,36) = 35.53, MSE = 2.62, p < 0.001*), and between ‘Phoneme’, ‘ROI’, and ‘Modality’ *(F(1,36) = 15.83, MSE = 3.54; p < 0.001)*. In subsequent 3-way ANOVAs split up by ‘Modality’, we observed significant 3-way interactions between ‘Phoneme’, ‘Area’ and ‘ROI’ in *<u>production</u> (F(1, 36) = 25.79, MSE = 3.34; p < 0.001)*, and in *<u>perception</u> (F(1,36) = 7.21, MSE = 2.66, p = 0.01).* Considering these significant interactions, next, and most crucially for the hypotheses, we performed 2-way ANOVAs split up by ‘Modality’ and ‘Area’ to assess whether the phoneme factor interacts with ROI across the language modalities and across the frontal and temporal cortices. In the *<u>motor cortex</u>* for *<u>production</u>* we observed a significant interaction between ‘Phoneme’ and ‘ROI’ *(F(1,36) = 56.31, MSE = 4.34, p < 0.001)*, indexing stronger activation of the Lip Motor Cortex when naming objects starting with bilabial speech sounds and stronger activation of the Tongue Motor Cortex when naming objects starting with alveolar speech sounds (***Fig. 2A*** and ***2B***). Importantly, also for *<u>perception</u>* the interaction between ‘Phoneme’ and ‘ROI’ was significant *(F(1,36) = 9.06, MSE = 2.43, p = 0.005),* likewise indicating that when participants passively listened to words starting with bilabial speech sounds there was more activity in the Lip Motor Cortex compared to words starting with alveolar speech sounds and vice versa in the Tongue Motor Cortex when listening to words starting with alveolar speech sounds (***Fig. 2A*** and ***2B***). Finally, when running 2-way ANOVAs in the *<u>temporal cortex</u>* there was no significant interaction between ‘Phoneme’ and ‘ROI’ in *<u>production</u> (F(1,36) = 2.59, MSE = 2.44, p = 0.12),* and no significant interaction between ‘Phoneme’ and ‘ROI’ in *<u>perception</u> (F(1,36) = 1.07, MSE = 2.11, p = 0.31)*, indicating similar activation profiles for bilabial and alveolar speech sounds across the modalities in the STG (***Fig. 2A*** and ***2B***); a result further confirmed by the absence of a significant interaction between ‘Phoneme’ and ‘Modality’ in the temporal cortex (*F(1,36) < 1*) (***Fig. 2A***). Instead, we observed a main effect of ‘Phoneme’ in the temporal cortex (*F(1,36) = 4.44, MSE = 7.52, p = 0.04*) (***Fig. 2A***), with stronger activations for words starting with alveolar speech sounds compared to bilabial speech sounds in both modalities and both regions of the STG (***Fig. 2B***).

**Fig. 2.**
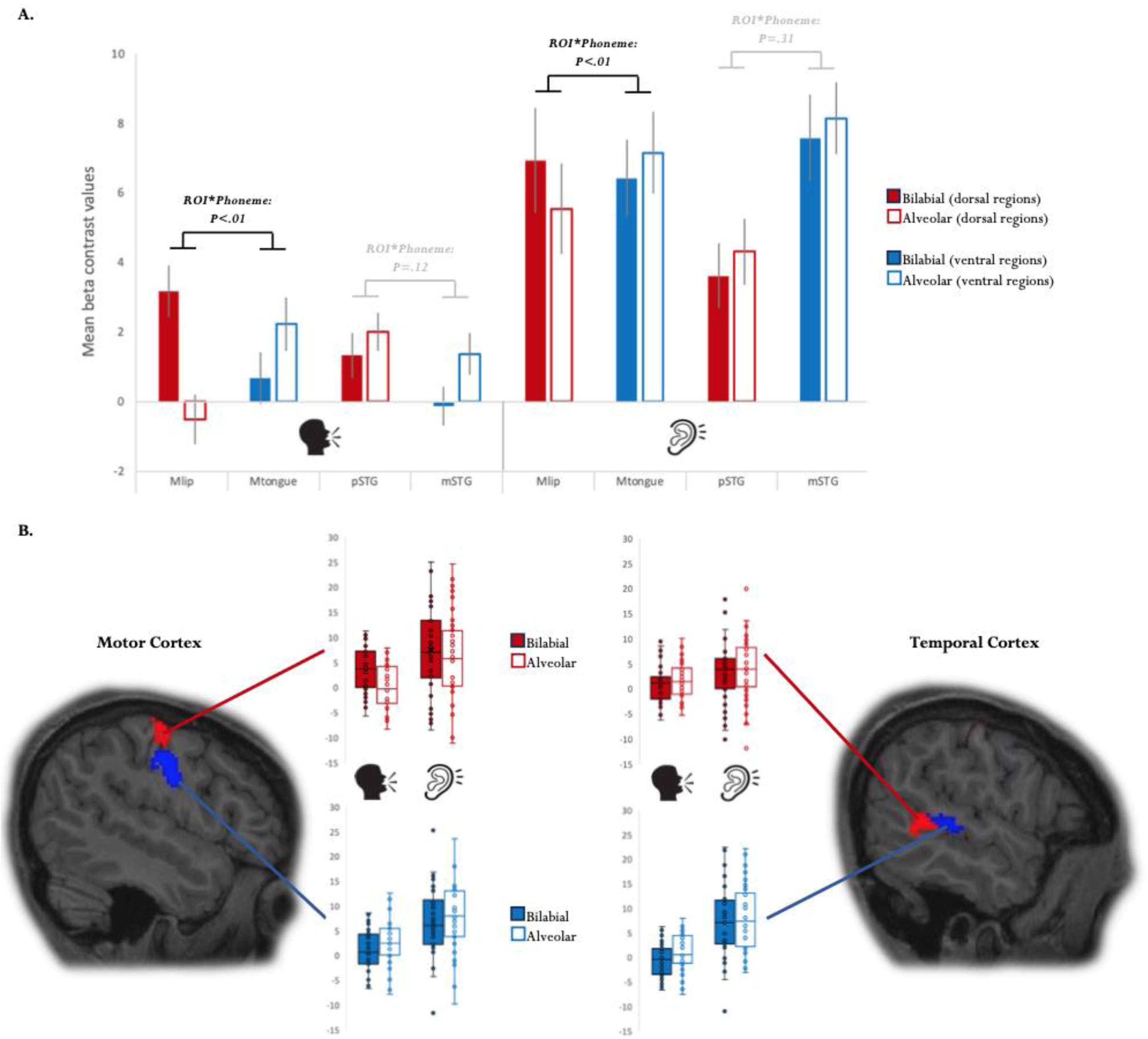
Results of the phoneme contrast between production and comprehension. **(A)** Mean voxel activation (with mean standard error) for bilabial-initial words (filled red and blue bars) and alveolar-initial words (outlined red and blue bars) in production (left) versus comprehension (right) for the different ROIs in the motor cortex (Mlip = lip-region in motor cortex; Mtongue = tongue-region in motor cortex) and temporal cortex (pSTG = posterior superior temporal gyrus; mSTG = middle superior temporal gyrus). The graph highlights the interactions between ROI and the Phoneme conditions which are identical across the language modalities: significant interactions in the motor cortex and the absence of significant interactions in the temporal cortex. **(B)** Dispersion graphs of the mean voxel activity of each participant for the different phoneme conditions (bilabial versus alveolar) in each of the four ROIs (motor regions left, temporal regions right) in production and comprehension. The graphs visualize the remarkable overlap between production and comprehension for the bilabial-alveolar contrast in all four ROIs. In the motor cortex, the red mask corresponds to the lip-region and the blue mask to the tongue-region. In the temporal cortex, the red mask corresponds to a more posterior portion of the superior temporal cortex and the blue mask to a more middle portion of the superior temporal cortex (note that the ROIs are based on individual masks).

## Discussion

In this fMRI study we set out to test whether the production and comprehension of words recruit phoneme-specific networks in the brain and whether those networks are shared across the language modalities. Thirty-seven participants performed both object naming and passive listening tasks on the same word stimuli. To trace cortical regions sensitive to phonological processing we used minimal pairs which solely differed in their initial phoneme: bilabial (e.g., ‘*ballon’*) or alveolar (e.g., ‘*talon’*). When comparing this phoneme contrast between the production and comprehension tasks, similar region by phoneme interactions were found in the motor cortex and the superior temporal cortex. In motor cortex a phoneme-specific topographical response was found for language production with bilabial words eliciting stronger activation in the motor region sensitive to lip movements and alveolar words eliciting stronger activation in the motor region sensitive to tongue movements (***Fig. 2***). Crucially, the phoneme by motor region interaction was also significant in the passive listening task, showing that within the same individuals those motor regions that responded in a phoneme-specific topographical manner during production, also responded in a phoneme-specific topographical manner during comprehension (***Fig. 2***). Furthermore, also in the superior temporal cortex we observed a similar activation pattern for the phoneme manipulation during object naming and passive listening: Both in more posterior as more middle portions of the STG alveolar-initial words produced stronger activations compared to bilabial-initial words (***Fig. 2***). Our study demonstrates a phoneme-specific fronto-temporal network when producing words which is identical to the network elicited when listening to words. This data pattern strongly supports the hypotheses of Integration Models (IM) (***Fig. 1***).

According to IM phonological representations of words will rely (at least) on temporal and frontal regions both when uttering as when listening to those words. This is because this class of models are based on assembly coding which allows to integrate correlated sensori-motor activation into neuronal circuits whose neurons are widely distributed across sensory, motor and convergence areas and strongly interlinked by long-distance cortico-cortical connections ^(59-61)^. When extending this neurobiological principle to cognition and language ^(27)^, co-processed auditory and motor properties of word forms become critical for interlinking perception- and production-related neurons in auditory and motor systems due to consistent correlated activation during acquisition ^(62,63)^. Therefore, in processing words, phonological information about the speech sounds making up the word form are processed in parallel in auditory brain regions in temporal cortex (acoustic-phonological features) and in motor-related brain regions in fronto-central cortex (articulatory-phonological features; ^(27,30)^). Recent evidence supports such parallel fronto-temporal phonological processing in speech production ^(31,33)^ and comprehension ^(28)^, along with its emergence in the first year of life ^(63)^. These results align with IM, as they assume shared mechanisms carrying both word production and perception. What differentiates between language modalities and tasks is the processing after these word representations have been activated, not the activation of the word representations itself ^(36)^. The current findings showing that both the production and comprehension of language elicits the same fronto-temporal brain responses that distinguish between and are specific to particular phonemic types perfectly fit the hypothesis of IM that phonological networks of words rely on shared neural representations across the language modalities.

Observing somatotopic motor activations when listening to speech is in line with previous fMRI findings ^(64-67; for a review see 68)^, but also extend these findings in important ways. Firstly, whereas previous neuroimaging work focused on the perception of meaningless syllables, we here demonstrate phoneme-specific motor activity elicited by meaningful words. Considering that words come with much greater phonetic variability compared to repeated syllables, observing phoneme-specific dissociations in motor cortex in the current set-up demonstrates robustness of the findings. It also addresses speculations that motor cortex involvement in speech comprehension might only be observable in artificial phonetic tasks, such as sound detection or classification, which are indeed not most relevant in day-to-day conversation ^(69,70)^. In contrast, producing or listening to meaningful language, as is the case here, concerns a more natural way of processing phonemes. A further noticeable extension of the current study compared to previous work is that both the comprehension and production of the same words were explored within the same brains. This leaves little doubt that the same motor regions recruited for the speech planning and articulation of the alveolar and bilabial minimal pairs are also recruited when passively listening to these words. Taken together, our study provides compelling evidence for the motor cortex’s involvement in language comprehension.

Both in posterior and middle portions of the superior temporal cortex, alveolar-initial words produced greater activation compared to bilabial-initial words during word production and comprehension. While, crucially, this activation pattern for phonological processing was again the same in production and comprehension, the pattern was different from the one found in the motor cortex in that the activation was not phoneme-specific in function of region within the STG. For IM we had suggested that similar region by phoneme interactions predicted for the motor cortex, may also be present in the temporal cortex. This suggestion had been based on data from previous work ^(42,44)^, and the theoretical consideration that if phonological networks include sensori-motor integration across production and perception, the spatial dissociations between alveolar and bilabial speech sounds in motor cortex may be communicated to temporal cortex in a topographic manner due to the general neighborhood-preserving nature of cortico-cortical connections ^(71)^. Categorical responses to different phonemic properties in the superior temporal cortex are well established ^(72-74)^. The absence of a dissociable superior-temporal brain response for the alveolar – bilabial contrast in the present study could be due to other acoustic features than place of articulation being more dominant for the organization of phonemes in the superior temporal cortex ^(48-50,74)^.

Despite the absence of topographic specificity of the brain response to the alveolar and bilabial stop consonants in superior temporal cortex, the activation again revealed the same pattern in language production and comprehension, with more activation for alveolar-initial words compared to bilabial-initial words across the modalities. Stronger neural responses to alveolar than bilabial speech sounds had previously been found in other production and perception studies^(42,44,45,73,75)^. This effect could be related to alveolar speech sounds having overall greater variability than bilabial speech sounds ^(76)^, and alveolar speech sounds being produced by a biomechanically more complex articulator (tongue) compared to bilabial speech sounds (lips) ^(42,75)^. Regardless of the exact reason why alveolar speech sounds activated STG more strongly than bilabial speech sounds, the global picture that emerges from our work is that the shared phonological network of words across the modalities is subserved by more general distributed representations in the superior temporal cortex linked to more phoneme-specific local topographies in motor cortex.

Integrating this overlapping spatial network observed for phonological processing with the fast parallel time-course of word processing observed across modalities ^(36)^, suggests that words rely on the same neural organization and activation dynamics during both language production and comprehension. This neural basis for words challenges traditional accounts and implies alternative mechanisms for differentiating production and comprehension in both healthy and brain-damaged individuals. One such mechanism could be that differences between production and comprehension arise not from distinct neural representations but from behavior-specific processing dynamics after word activation ^(31,36,77)^. The memory representation of a word would activate similarly during speaking and listening, but the subsequent goal-directed processing, such as articulating the word or integrating it in the prior context, would vary by language behavior. This framework, which combines shared memory representations across the language modalities with distinct goal-directed processing, may offer novel perspectives to explain important production-perception phenomena in the cognitive and neurosciences such as neuropsychological double dissociations between production and comprehension ^(78)^, how words are integrated across the modalities to achieve combinatorial processing ^(79)^, how to engage in dialogue ^(80)^, and how we can monitor our speech for errors ^(81)^.

## Conclusion

In this fMRI study with 37 participants, we asked whether phoneme-specific networks can be observed in both language production and comprehension. We found for both language modalities the same pattern of haeamodynamic brain activity specific to different phoneme types: Minimal pairs differing only in the word-initial alveolar or bilabial phoneme activated the motor cortex topographically and the superior temporal cortex to different degrees, regardless of whether participants were uttering or listening to those minimal pairs. We argue that this pattern meets the predictions of Integration Models ^(27-33)^, according to which the cortical representations of words are neuronal circuits that are shared between language production and comprehension. The results indicate that language representations are not constrained by a strict sensory-motor division. Instead, our data suggest that language comprehension can involve motor areas of the brain, and language production can involve perceptual areas of the brain. And while phonology in the brain clearly encompasses more than the prediction-specific regions focused on here, our findings show that within the same individual phoneme-specific motor topography during speaking is also active during listening, and more general phoneme-sensitive superior temporal activity during listening manifests also when speaking. This is a particularly compelling test of the notion that word representations, including their phonological properties, are integrated across and general to both language modalities.

## Materials & Methods

We have made the code, analyses pipelines, materials, averaged data and all statistical analyses of the processed data available in the project OSF repository at https://osf.io/bhfdw/ (the raw, preprocessed and structural MRI data, which are too large in size for the OSF repository, will be made available upon request to the corresponding author).

### Participants

Forty-four native French speakers with normal hearing, normal or corrected eyesight, and no history of any neurological problems participated in the study. We excluded 7 participants from the study due to technical issues, resulting in a total of 37 participants (female = 27, median age = 25.5, range = 18.5 - 36.2). All of the participants were recruited on the voluntary basis through the Aix-Marseille University mailing lists. The study received ethical approval (filed under Id 2017-A03614-49 from the regional ethical committee, Comité de protection des personnes sud Méditerranée I). All participants gave written consent to participate in the study and received monetary compensation for their participation.

### Stimuli

Twenty minimal phonological pairs of nouns were selected from the Lexique database on French language ^(52)^. Selection criteria included that minimal word pairs could only differ in their starting consonant, namely, bilabial */b/* or */p/* for one member of the minimal pair, and an alveolar consonant, */d/* or */t/*, for the other member. Items were controlled for length (1-3 syllables), lexical frequency, and had to be concrete, depictable nouns (since the same stimuli were presented in auditory and visual modality; see below). Analyses of the naming latencies in the production task (standard picture naming) furthermore confirmed that the items were indeed well-controlled since no differences in reaction times were found between the bilabial words (783ms) and alveolar words (785ms) (details as well as all individual naming latencies for all items can be found in the online repository: https://osf.io/bhfdw/). An additional set of 10 filler nouns and 5 practice nouns were selected which, unlike the target pairs, did not start with alveolar or bilabial consonants. All stimuli were presented in the visual modality for the production task (object naming) and the auditory modality for the perception task (passive listening) (see https://osf.io/bhfdw/ for an overview of all stimuli).

For the visual modality, 40 grey-scale PNG images depicting the minimal pairs (targets), as well as 10 filler images, and 5 practice images were selected. Out of those, 37 images (22 targets, 10 filler items, and 5 practice items) were selected from the MultiPic database ^(53)^, and an additional 18 target images were selected from the open-source grey-scale images database Google Line Drawings. All images were matched in pixel-size.

For the auditory modality, the target, filler and practice stimuli were pre-recorded by a native female French speaker in the soundproof room of the Laboratoire Parole et Langage at Aix-Marseille University using an RME fireface UC audio interface and a headset Cardioid Condenser Microphone (AKG C520) at a sampling rate of 44 100 Hz. The tokens were preprocessed and denoised via AudaCity® software. Average duration of audio stimuli was 640 ms (range: 304-841 ms), and there were no differences in average length between the bilabial and alveolar stimuli sets.

Prior to the experiment (and outside the scanner) the participants were familiarized with all target and filler items by presenting randomly each object accompanied with its corresponding auditory name. This was done to substantially reduce potential naming errors and potential repetition effects during the main experiment (since errors and repetition effects are drastically reduced for objects and words after the first repetition ^(38)^). During the main experiment, each target and filler appeared eight times for each participant (four times in the production task and four times in the perception task), leading to a total of 400 trials (160 target trials + 40 filler trials in production and 160 target trials + 40 filler trials in perception). In this manner, for each task, there were four runs (40 targets + 10 fillers), which were semi-randomized for each run following the constraint that the same starting consonant (/b/, /p/, /d/ and /t/) could not be presented in sequence for more than 2 times.

### Tasks and Procedure

The main experiment consisted of two tasks, a standard production task and a standard perception task (***Fig. 1***).

*The production task* was object naming, where participants are presented with an object picture for which they have to utter its most typical designation or ‘name’ as rapidly and correctly as possible. Trial structure was the following (***Fig. 1***, *upper panel*): A fixation cross was presented in the center with variable duration between trials (randomly varying between 500 – 1200 ms to create inter-stimulus jitter). This was followed by object presentation for 640 ms (which corresponds to the average duration of the auditory stimuli in the perception task to ensure that overall target duration was the same between the production and perception tasks). Object presentation was followed by a blank screen presented for 2500 ms.

*The perception task* was a go/no-go paradigm where the no-go trials corresponded to the auditory presentation of the target stimuli (passive listening; 80% of the trials), and the go-trials included the auditory presentation of the filler stimuli followed by two alternative pictures (20% of the trials) (***Fig. 1****, lower panel*). On those go-trials participants had to perform word-to-picture matching to ensure semantic processing of the auditory input: the two images following the filler word appeared on the left and right of the screen and participants had to push either a left or right response button to indicate which object corresponded to the just heard word. Participants had to press the response buttons with their left middle and index fingers so as to minimize left-hemispheric response-related cortical activity. All images were chosen from the pool of filler stimuli, and response fingers were fully counterbalanced across participants. For analyses, only the no-go trials (passive listening) are analyzed. Trial structure of the no-go trials fully matched that of the production task (***Fig. 1***): A trial started with jittered fixation cross presentation (between 500 – 1200 ms), followed by the audio stimulus which was presented on average for 640 ms (range: 304 and 841 ms), and ending with a blank screen during 2500 ms. In the case of the filler trials, instead of a blank screen after the audio stimulus the participants were presented with two images for 2500 ms during which the word-to-picture decision could be made. For each run a baseline period of 5040 ms at the beginning and 12000 ms at the end was introduced.

### MRI Data Acquisition

Functional images were acquired at the Marseille MRI Center, using a 3-T Siemens Magnetom Prisma MR system with a 64-channel head coil. Functional images covering the whole brain were acquired during task performance, using a multiband blood oxygen level dependent (BOLD)-sensitive gradient echo planar imaging (EPI) sequence ^(54)^ (repetition time [TR] = 1040 ms, echo time [TE] = 35.2 ms, flip angle = 62°, 65 slices, field of view [FOV] = 200 × 200 mm2, matrix = 100 × 100, slice thickness = 2 mm, multiband factor = 5, bandwidth = 2380 Hz/pixel). Whole-brain anatomical magnetic resonance imaging (MRI) data were acquired using high-resolution structural T1-weighted image (MPRAGE sequence, voxel size = 1 × 1 × 1 mm3, data matrix 256 x 240 x 192, TR/TI (inversion time)/TE = 2300/925/2.98 ms, flip angle = 9°, bandwidth = 240 Hz/pixel, GRAPPA = 3). Prior to functional imaging and to correct functional images for susceptibility induced distortions, a pair of spin-echo EPI sequences was acquired twice with opposite phase encode directions along the anterior-posterior axis with the following parameters: TR/TE = 7106/58 ms, voxel size = 2 × 2 × 2 mm3, slices = 65, FOV = 200 × 200 mm2. Participants’ movements during data acquisition were controlled using Framewise Integrated Real Time MRI Monitoring ^(55)^.

### Image processing and analyses

MRI data were preprocessing performed using *fMRIPrep* 20.2.2 (RRID:SCR_016216)^1^, which is based on *Nipype* 1.6.1 (RRID:SCR_002502)^2^.

### Anatomical data preprocessing

The T1-weighted (T1w) image was corrected for intensity non-uniformity (INU) with N4BiasFieldCorrection ^(56)^, distributed with ANTs 2.3.3 (RRID:SCR_004757)^3^, and used as T1w-reference throughout the workflow. The T1w-reference was then skull-stripped with a *Nipype* implementation of the antsBrainExtraction.sh workflow (from ANTs), using OASIS30ANTs as target template. Brain tissue segmentation of cerebrospinal fluid (CSF), white-matter (WM) and grey-matter (GM) was performed on the brain-extracted T1w using fast (FSL 5.0.9, RRID:SCR_002823)^4^. Volume-based spatial normalisation to one standard space (MNI152NLin2009cAsym) was performed through nonlinear registration with antsRegistration (ANTs 2.3.3), using brain-extracted versions of both T1w reference and the T1w template. The following template was selected for spatial normalisation: *ICBM 152 Nonlinear Asymmetrical template version 2009c* (RRID:SCR_008796; TemplateFlow ID: MNI152NLin2009cAsym)^5^.

### Functional data preprocessing

For each of the 16 BOLD runs found per subject (2 experimental tasks and 2 localizers), the following preprocessing was performed. First, a reference volume and its skull-stripped version were generated by aligning and averaging 1 single-band references. Head-motion parameters with respect to the BOLD reference (transformation matrices, and six corresponding rotation and translation parameters) are estimated before any spatiotemporal filtering using mcflirt (FSL 5.0.9)^6^. A B0-nonuniformity map (or *fieldmap*) was estimated based on two (or more) echo-planar imaging (EPI) references with opposing phase-encoding directions, with 3dQwarp Cox and Hyde (AFNI 20160207). Based on the estimated susceptibility distortion, a corrected EPI (echo-planar imaging) reference was calculated for a more accurate co-registration with the anatomical reference. The BOLD time-series (including slice-timing correction when applied) were resampled onto their original, native space by applying a single, composite transform to correct for head-motion and susceptibility distortions. These resampled BOLD time-series will be referred to as *preprocessed BOLD in original space*, or just *preprocessed BOLD*. The BOLD reference was then co-registered to the T1w reference using flirt (FSL 5.0.9)^7^ with the boundary-based registration cost-function^8^. Co-registration was configured with nine degrees of freedom to account for distortions remaining in the BOLD reference. First, a reference volume and its skull-stripped version were generated using a custom methodology of *fMRIPrep*. Several confounding time-series were calculated based on the *preprocessed BOLD*: framewise displacement (FD), DVARS and three region-wise global signals. FD was computed using two formulations following Power (absolute sum of relative motions)^9^ and Jenkinson (relative root mean square displacement between affines)^10^. FD and DVARS are calculated for each functional run, both using their implementations in *Nipype* (following the definitions by Power et al. 2014). The three global signals are extracted within the CSF, the WM, and the whole-brain masks. Additionally, a set of physiological regressors were extracted to allow for component-based noise correction (*CompCor*)^11^. Principal components are estimated after high-pass filtering the *preprocessed BOLD* time-series (using a discrete cosine filter with 128s cut-off) for the two *CompCor* variants: temporal (tCompCor) and anatomical (aCompCor). tCompCor components are then calculated from the top 2% variable voxels within the brain mask. For aCompCor, three probabilistic masks (CSF, WM and combined CSF+WM) are generated in anatomical space. The implementation differs from that of Behzadi et al. in that instead of eroding the masks by 2 pixels on BOLD space, the aCompCor masks are subtracted a mask of pixels that likely contain a volume fraction of GM. This mask is obtained by thresholding the corresponding partial volume map at 0.05, and it ensures components are not extracted from voxels containing a minimal fraction of GM. Finally, these masks are resampled into BOLD space and binarized by thresholding at 0.99 (as in the original implementation). Components are also calculated separately within the WM and CSF masks. For each CompCor decomposition, the *k* components with the largest singular values are retained, such that the retained components’ time series are sufficient to explain 50 percent of variance across the nuisance mask (CSF, WM, combined, or temporal). The remaining components are dropped from consideration. The head-motion estimates calculated in the correction step were also placed within the corresponding confounds file. The confound time series derived from head motion estimates and global signals were expanded with the inclusion of temporal derivatives and quadratic terms for each^12^. Frames that exceeded a threshold of 0.5 mm FD or 2.0 standardised DVARS were annotated as motion outliers. The BOLD time-series were resampled into standard space, generating a *preprocessed BOLD run in MNI152NLin2009cAsym space*. First, a reference volume and its skull-stripped version were generated using a custom methodology of *fMRIPrep*. All resamplings can be performed with *a single interpolation step* by composing all the pertinent transformations (i.e. head-motion transform matrices, susceptibility distortion correction when available, and co-registrations to anatomical and output spaces). Gridded (volumetric) resamplings were performed using antsApplyTransforms (ANTs), configured with Lanczos interpolation to minimise the smoothing effects of other kernels. Non-gridded (surface) resamplings were performed using mri_vol2surf (FreeSurfer). Many internal operations of *fMRIPrep* use *Nilearn* 0.6.2 (RRID:SCR_001362)^13^, mostly within the functional processing workflow.

### Functional localizer tasks

At the end of each session 2 functional localizer tasks using a blocked design were performed in counterbalanced order. One of the localizer tasks was designed to assess within each individual the spatial dissociation in the motor cortex between lip movements and tongue movements (based on the lip-tongue motor localizer in ^(44)^). This was done to obtain functional regions of interest (ROIs) for the bilabial (lip-related) vs alveolar (tongue-related) conditions in the main experiment. Each of the two runs consisted of eight blocks of 16 seconds each; during one block the word *“levres” (lips)* was presented on the screen, while during the other block the word *“langue” (tongue)* was presented on the screen. Participants were instructed to slightly move their lips for the entire duration that the word *“levres”* was presented, and slightly tap their tongue on the palate for the entire duration the word *“langue”* was presented. No breaks were introduced between the blocks, and baselines of 5040 ms and 13040 ms were recorded at the beginning and end of each run respectively.

In the other localizer task participants listened to syllables which either started with a bilabial or alveolar speech sound to define individual specific functional ROIs underlying the phonological processing of bilabial and alveolar consonants in the superior temporal gyrus. Each of the two runs consisted of 8 blocks of bilabial-initial *(/pa/, /ba/, /ma/)* or alveolar-initial *(/da/, /ta/, /sa/, /za/, /na/)* syllables, and each block lasted for 16 s.

### Analyses

The preprocessed data were imported to and analyzed using the Statistical Parametric Mapping software (SPM12; https://www.Fil.ion.ucl.ac.uk/spm/software/spm12/) in MATLAB R2018b (Mathworks Inc., Natick, MA). Smoothing was performed with an isotropic Gaussian kernel (full-width at half-maximum = 5 mm). For the univariate ROIs analyses a General Linear Model (GLM) was built for each participant for each experimental task (production and perception), as well as for the motor and syllable localizers tasks. Regressors of no interest included 24 head movement regressors, the mean signal of the CSF and WM mask, and 24 aCompCor regressors related to CSF and WM.

For the motor localizer task, the GLM included, for each of the 4 runs, the following regressors of interest: ‘Lip-movement’ and ‘Tongue-movement’. To calculate the functional ROIs corresponding to the lips and the tongue within the motor cortex, individual maps were extracted based on the GLM output (contrasts: ‘Lip-movement’ vs ‘baseline’, ‘Tongue-movement’ vs ‘baseline‘). In addition, to avoid including non-motor related activity, we added an anatomical constraint to only consider activity within the sensorimotor cortex (based on the 400-parcels atlas of Schaefer ^(57)^). In this manner, our functional ROIs concerned masks of the intersection between functional and anatomical constraints within each individual (***Fig. 2***).

For the auditory syllable localizers, the GLM did not result in any reliable ROIs within the temporal cortex^14^. Therefore, we defined the ROIs in the temporal cortex based on the intersection of the individual grey matter with pre-defined anatomical regions from the 400-parcel atlas of Schaefer ^(57)^. Concretely, based on the previous literature ^(42,44)^ we selected two anatomical parcels in the temporal cortex corresponding to the middle portion of the STG (parcel 197) and the posterior portion of the STG (parcel 198).

For the ROI analyses of the main experiment, GLMs were created for each of the 8 runs and each of the 4 ROIs identified above, which contained the following regressors of interest: ‘bilabial_production’, ‘alveolar_production’, ‘bilabial_perception, ‘alveolar_perception’, ‘filler_production’, and ‘filler_perception’. Within each of the four ROI masks we extracted for each of the regressors the mean voxel activation within each participant. For statistical analyses the contrast values of the experimental variables were created by subtracting the experimental regressors from the filler regressors (i.e., bilabial_production – filler_production; alveolar_production – filler_production; bilabial_perception – filler_perception; alveolar_perception – filler_perception). Analysis of Variance (ANOVA) were run starting with all experimental variables, and subsequently breaking down those variables which displayed significant interactions. In this manner, the highest-level analysis concerned a 4-way 2x2x2x2 ANOVA including the following variables (with the corresponding levels in brackets): ‘Phoneme’ (bilabial vs. alveolar) x ‘Area’ (motor cortex vs. superior temporal cortex) x ‘Region of Interest (ROI)’ (anterior (tongue-related) vs posterior (lip-related)) x ‘Modality’ (production vs. comprehension).

## Acknowledgements

We are grateful to numerous colleagues of the LPL and the INT at Aix-Marseille University and the Brain Language Laboratory at the Free University Berlin for the discussions and comments on the results and their significance. K.S. was supported by the ‘PhoNet’ grant of the “Agence National de la Recherche” (ANR) to conduct this study (ANR-18-FRAL-0013-01). Furthermore, the study received additional support by an ANR grant awarded to the Institute of Language, Communication and Brain (ANR-16-CONV-0002).

## Author contributions

K.S. conceptualized and supervised the study. X.D. and K.S. designed the study. X.D., J-L.A., E.R., J.S., B.N., S.D., F.P. and K.S. implemented the experimental and functional recording set-up of the study. X.D., A.F. and B.I. collected the data. X.D., J-L.A., E.R. and K.S. analyzed the data. X.D. and K.S. interpreted the data and wrote the initial draft. All authors reviewed and commented on the manuscript.

## Footnotes

1 Esteban O, et al., 2018.

2 Gorgolewski et al., 2011; 2018.

3 Avants et al., 2008.

4 Zhang, et al., 2001.

5 Fonov et al., 2009.

6 Jenkinson et al., 2002.

7 Jenkinson and Smith, 2001.

8 Greve and Fischl, 2009.

9 Power et al., 2014.

10 Jenkinson et al., 2002.

11 Behzadi et al., 2007.

12 Satterthwaite et al., 2013.

13 Abraham et al., 2014.

14 Note that for the localizers we tried both the standard individual functional ROIs approach as well as the group-constrained subject-specific approach developed by Federenko and colleagues ^(58)^. While for the motor localizer results were comparable, for the auditory localizer neither approach produced significant ROIs in the temporal cortex; indicating that our auditory syllable task is likely not a good localizer task to highlight differences in phoneme processing within the temporal cortex.

## Notes

https://osf.io/bhfdw/

### Competing Interest Statement

The authors have declared no competing interest.

### Summary of Updates

This revised version contains the following changes compared to the previous version: - Inclusion and discussion of a few more studies, in particular ECoG studies - Inclusion of naming latencies for the production experiment and made available all individual audio files of each naming response.

## References

1. Pickering, M. J., & Garrod, S. An integrated theory of language production and comprehension. Behav. Brain Sci., 36(4), 329–347 (2013).

2. Levinson, S. C. Turn-taking in human communication–origins and implications for language processing. Trends Cogn. Sci., 20(1), 6–14 (2016).

3. Geschwind, N. Specializations of the human brain. Sci. Am., 241(3), 180–201 (1979).

4. Damasio, A. R., & Geschwind, N. The neural basis of language. Annu. Rev. Neurosci., 7(1), 127–147 (1984).

5. Price, C. J. A review and synthesis of the first 20 years of PET and fMRI studies of heard speech, spoken language and reading. Neuroimage, 62(2), 816–847 (2012).

6. Caplan, D. Neurolinguistics and linguistic aphasiology: An introduction. Cambridge University Press (1987).

7. Damasio, H., & Damasio, A. R. The anatomical basis of conduction aphasia. Brain, 103(2), 337–350 (1980).

8. Caramazza, A., & Zurif, E. B. Dissociation of algorithmic and heuristic processes in language comprehension: Evidence from aphasia. Brain Lang., 3(4), 572–582 (1976).

9. Miceli, G., & Caramazza, A. Dissociation of inflectional and derivational morphology. Brain Lang., 35(1), 24–65 (1988).

10. Dreyer, F. R., Picht, T., Frey, D., Vajkoczy, P., & Pulvermüller, F. The functional relevance of dorsal motor systems for processing tool nouns–evidence from patients with focal lesions. Neuropsychologia, 141, 107384 (2020).

11. Dell, G. S., & O’Seaghdha, P. G. Stages of lexical access in language production. Cognition, 42(1-3), 287–314 (1992).

12. Levelt, W. J., Roelofs, A., & Meyer, A. S. A theory of lexical access in speech production. Behav. Brain Sci., 22(1), 1–38 (1999).

13. Meyer, A. S., Huettig, F., & Levelt, W. J. Same, different, or closely related: What is the relationship between language production and comprehension?. J. Mem. Lang., 89, 1–7 (2016).

14. Stephens, G. J., Silbert, L. J., & Hasson, U. Speaker–listener neural coupling underlies successful communication. Proc. Natl. Acad. Sci. U.S.A., 107(32), 14425–14430 (2010).

15. Segaert, K., Menenti, L., Weber, K., Petersson, K. M., & Hagoort, P. Shared syntax in language production and language comprehension—an fMRI study. Cereb. Cortex, 22(7), 1662–1670 (2012).

16. Dikker, S., Silbert, L. J., Hasson, U., & Zevin, J. D. On the same wavelength: predictable language enhances speaker–listener brain-to-brain synchrony in posterior superior temporal gyrus. J. Neurosci., 34(18), 6267–6272 (2014).

17. Silbert, L. J., Honey, C. J., Simony, E., Poeppel, D., & Hasson, U. Coupled neural systems underlie the production and comprehension of naturalistic narrative speech. Proc. Natl. Acad. Sci. U.S.A., 111(43), E4687–E4696 (2014).

18. Giglio, L., Ostarek, M., Weber, K., & Hagoort, P. Commonalities and asymmetries in the neurobiological infrastructure for language production and comprehension. Cereb. Cortex, 32(7), 1405–1418 (2022).

19. Giglio, L., Ostarek, M., Sharoh, D., & Hagoort, P. Diverging neural dynamics for syntactic structure building in naturalistic speaking and listening. Proc. Natl. Acad. Sci. U.S.A., 121(11), e2310766121 (2024).

20. Matchin, W., & Wood, E. Syntax-sensitive regions of the posterior inferior frontal gyrus and the posterior temporal lobe are differentially recruited by production and perception. Cereb. Cortex Commun., 1(1), tgaa029 (2020).

21. Indefrey, P., & Levelt, W. J. The spatial and temporal signatures of word production components. Cognition, 92(1-2), 101–144 (2004).

22. Hickok, G., & Poeppel, D. The cortical organization of speech processing. Nat. Rev. Neurosci., 8(5), 393–402 (2007).

23. Indefrey, P. The spatial and temporal signatures of word production components: a critical update. Front. Psychol., 2, 255 (2011).

24. Hickok, G. The architecture of speech production and the role of the phoneme in speech processing. Lang. Cogn. Neurosci., 29(1), 2–20 (2014).

25. Roelofs, A. A dorsal-pathway account of aphasic language production: The WEAVER++/ARC model. Cortex, 59, 33–48 (2014).

26. de Zubicaray, G. I. The Neural Organization of Language Production: Evidence from Neuroimaging and Neuromodulation. Lang. Prod., 97–142 (2023).

27. Pulvermüller, F. Words in the brain’s language. Behav. Brain Sci., 22(2), 253–279 (1999).

28. Pulvermüller, F. Neural reuse of action perception circuits for language, concepts and communication. Prog. Neurobiol., 160, 1–44 (2018).

29. Tourville, J. A., & Guenther, F. H. The DIVA model: A neural theory of speech acquisition and production. Lang. Cogn. Process., 26(7), 952–981 (2011).

30. Pulvermüller, F., & Fadiga, L. Active perception: sensorimotor circuits as a cortical basis for language. Nat. Rev. Neurosci., 11(5), 351–360 (2010).

31. Strijkers, K., & Costa, A. The cortical dynamics of speaking: Present shortcomings and future avenues. Lang. Cogn. Neurosci., 31(4), 484–503 (2016).

32. Strijkers, K. A neural assembly–based view on word production: The bilingual test case. Lang. Learn., 66(S2), 92–131 (2016).

33. Kerr, E., Ivanova, B., & Strijkers, K. Lexical access in speech production: Psycho-and neurolinguistic perspectives on the spatiotemporal dynamics. In Lang. Prod. (pp. 32–65) Routledge (2023).

34. Okada, K., & Hickok, G. Left posterior auditory-related cortices participate both in speech perception and speech production: Neural overlap revealed by fMRI. Brain Lang., 98(1), 112–117 (2006).

35. Strijkers, K., & Costa, A. Riding the lexical speedway: A critical review on the time course of lexical selection in speech production. Front. Psychol., 2, 14253 (2011).

36. Fairs, A., Michelas, A., Dufour, S., & Strijkers, K. The same ultra-rapid parallel brain dynamics underpin the production and perception of speech. Cereb. Cortex Commun., 2(3), tgab040 (2021).

37. Vitevitch, M. S., & Luce, P. A. Probabilistic phonotactics and neighborhood activation in spoken word recognition. J. Mem. Lang., 40(3), 374–408 (1999).

38. Strijkers, K., Costa, A., & Thierry, G. Tracking lexical access in speech production: electrophysiological correlates of word frequency and cognate effects. Cereb. Cortex, 20(4), 912–928 (2010).

39. Guenther, F. H., Ghosh, S. S., & Tourville, J. A. Neural modeling and imaging of the cortical interactions underlying syllable production. Brain Lang., 96(3), 280–301 (2006).

40. Bouchard, K. E., Mesgarani, N., Johnson, K., & Chang, E. F. Functional organization of human sensorimotor cortex for speech articulation. Nature, 495(7441), 327–332 (2013).

41. Murakami, T., Kell, C. A., Restle, J., Ugawa, Y., & Ziemann, U. Left dorsal speech stream components and their contribution to phonological processing. J. Neurosci., 35(4), 1411–1422 (2015).

42. Strijkers, K., Costa, A., & Pulvermüller, F. The cortical dynamics of speaking: Lexical and phonological knowledge simultaneously recruit the frontal and temporal cortex within 200 ms. NeuroImage, 163, 206–219 (2017).

43. Fadiga, L., Craighero, L., Buccino, G., & Rizzolatti, G. Speech listening specifically modulates the excitability of tongue muscles: a TMS study. Eur. J. Neurosci., 15(2), 399–402 (2002).

44. Pulvermüller, F., Huss, M., Kherif, F., Moscoso del Prado Martin, F., Hauk, O., & Shtyrov, Y. Motor cortex maps articulatory features of speech sounds. Proc. Natl. Acad. Sci., 103(20), 7865–7870 (2006).

45. D’Ausilio, A., Pulvermüller, F., Salmas, P., Bufalari, I., Begliomini, C., & Fadiga, L. The motor somatotopy of speech perception. Curr. Biol., 19(5), 381–385 (2009).

46. Möttönen, R., & Watkins, K. E. Motor representations of articulators contribute to categorical perception of speech sounds. J. Neurosci., 29(31), 9819–9825 (2009).

47. Schomers, M. R., Kirilina, E., Weigand, A., Bajbouj, M., & Pulvermüller, F. Causal influence of articulatory motor cortex on comprehending single spoken words: TMS evidence. Cereb. Cortex, 25(10), 3894–3902 (2015).

48. Mesgarani, N., Cheung, C., Johnson, K., & Chang, E. F. Phonetic feature encoding in human superior temporal gyrus. Science, 343(6174), 1006–1010 (2014).

49. Bhaya-Grossman, I., & Chang, E. F. Speech computations of the human superior temporal gyrus. Annu. Rev. Psychol., 73(1), 79–102 (2022).

50. Cheung, C., Hamilton, L. S., Johnson, K., & Chang, E. F. The auditory representation of speech sounds in human motor cortex. Elife, 5, e12577 (2016).

51. Skipper, J. I., Devlin, J. T., & Lametti, D. R. The hearing ear is always found close to the speaking tongue: Review of the role of the motor system in speech perception. Brain Lang., 164, 77–105 (2017).

52. New, B., Pallier, C., Brysbaert, M., & Ferrand, L. Lexique 2: A new French lexical database. Behav. Res. Methods Instrum. Comput., 36(3), 516–524 (2004).

53. Duñabeitia, J. A., Crepaldi, D., Meyer, A. S., New, B., Pliatsikas, C., Smolka, E., & Brysbaert, M. MultiPic: A standardized set of 750 drawings with norms for six European languages. Q. J. Exp. Psychol., 71(4), 808–816 (2018).

54. Moeller, S., Yacoub, E., Olman, C. A., Auerbach, E., Strupp, J., Harel, N., & Uğurbil, K. Multiband multislice GE-EPI at 7 tesla, with 16-fold acceleration using partial parallel imaging with application to high spatial and temporal whole-brain fMRI. Magn. Reson. Med., 63(5), 1144–1153 (2010).

55. Dosenbach, N. U., Koller, J. M., Earl, E. A., Miranda-Dominguez, O., Klein, R. L., Van, A. N., … & Fair, D. A. Real-time motion analytics during brain MRI improve data quality and reduce costs. NeuroImage, 161, 80–93 (2017).

56. Tustison, N. J., Avants, B. B., Cook, P. A., Zheng, Y., Egan, A., Yushkevich, P. A., & Gee, J. C. N4ITK: improved N3 bias correction. IEEE Trans. Med. Imaging, 29(6), 1310–1320 (2010).

57. Schaefer, A., Kong, R., Gordon, E. M., Laumann, T. O., Zuo, X. N., Holmes, A. J., … & Yeo, B.T. Local-global parcellation of the human cerebral cortex from intrinsic functional connectivity MRI. Cereb. Cortex, 28(9), 3095–3114 (2018).

58. Fedorenko, E., Hsieh, P. J., Nieto-Castañón, A., Whitfield-Gabrieli, S., & Kanwisher, N. New method for fMRI investigations of language: defining ROIs functionally in individual subjects. J. Neurophysiol., 104(2), 1177–1194 (2010).

59. Hebb, D.O. The Organization of Behaviour. A Neuropsychological Theory. John Wiley, New York. (1949).

60. Braitenberg, V. Cell assemblies in the cerebral cortex. In Heim, R. & Palm, G. (Eds), Theoretical Approaches to Complex Systems, Lecture Notes in Biomathematics, Vol. 21. Springer Verlag, Berlin, pp. 171–188 (1978).

61. Artola, A. & Singer, W. Long-term depression of excitatory synaptic transmission and its relationship to long-term potentiation. Trends Neurosci., 16, 480–487 (1993).

62. Werker, J. F., & Tees, R. C. Influences on infant speech processing: Toward a new synthesis. Annu. Rev. Psychol., 50(1), 509–535 (1999).

63. Kuhl, P. K. Brain mechanisms in early language acquisition. Neuron, 67(5), 713–727 (2010).

64. Wilson, S. M., Saygin, A. P., Sereno, M. I., & Iacoboni, M. Listening to speech activates motor areas involved in speech production. Nat. Neurosci., 7(7), 701–702 (2004).

65. Skipper, J. I., Goldin-Meadow, S., Nusbaum, H. C., & Small, S. L. Speech-associated gestures, Broca’s area, and the human mirror system. Brain Lang., 101(3), 260–277 (2007).

66. Correia, J. M., Jansma, B. M., & Bonte, M. Decoding articulatory features from fMRI responses in dorsal speech regions. J. Neurosci., 35(45), 15015–15025 (2015).

67. Evans, S., & Davis, M. H. Hierarchical organization of auditory and motor representations in speech perception: evidence from searchlight similarity analysis. Cereb. Cortex, 25(12), 4772–4788 (2015).

68. Schomers, M. R., & Pulvermüller, F. Is the sensorimotor cortex relevant for speech perception and understanding? An integrative review. Front. Hum. Neurosci., 10, 435 (2016).

69. Mahon, B. Z., & Caramazza, A. A critical look at the embodied cognition hypothesis and a new proposal for grounding conceptual content. J. Physiol.-Paris, 102(1-3), 59–70 (2008).

70. Hickok, G. The myth of mirror neurons: The real neuroscience of communication and cognition. WW Norton & Company. (2014).

71. Braitenberg, V., & Schuz, A. *Cortex: Statistics and Geometry of Neuronal Connectivity*. Springer, Berlin. (1998).

72. Obleser, J., Elbert, T., Lahiri, A., & Eulitz, C. Cortical representation of vowels reflects acoustic dissimilarity determined by formant frequencies. *Cogn*. Brain Res., 15(3), 207–213 (2003).

73. Chang, E. F., Rieger, J. W., Johnson, K., Berger, M. S., Barbaro, N. M., & Knight, R. T. Categorical speech representation in human superior temporal gyrus. Nat. Neurosci., 13(11), 1428–1432 (2010).

74. Arsenault, J. S., & Buchsbaum, B. R. Distributed neural representations of phonological features during speech perception. J. Neurosci., 35(2), 634–642 (2015).

75. Bartoli, E., D’Ausilio, A., Berry, J., Badino, L., Bever, T., & Fadiga, L. Listener–speaker perceived distance predicts the degree of motor contribution to speech perception. Cereb. Cortex, 25(2), 281–288 (2015).

76. Chomsky, N., & Halle, M. The sound pattern of English. Harper & Row, New York. (1968).

77. Pulvermüller, F. The neuroscience of language: On brain circuits of words and serial order. Cambridge University Press (2002).

78. Shallice, T. From neuropsychology to mental structure. Cambridge University Press (1988).

79. Hagoort, P., & Indefrey, P. The neurobiology of language beyond single words. Annu. Rev. Neurosci., 37(1), 347–362 (2014).

80. Pickering, M. J., & Garrod, S. Understanding dialogue: Language use and social interaction. Cambridge University Press. (2021).

81. Runnqvist, E., Chanoine, V., Strijkers, K., Pattamadilok, C., Bonnard, M., Nazarian, B., … & Alario, F.X. Cerebellar and cortical correlates of internal and external speech error monitoring. Cereb. Cortex Commun., 2(2), tgab038 (2021).

